# Feature Specific Prediction Errors and Surprise across Macaque Fronto-Striatal Circuits during Attention and Learning

**DOI:** 10.1101/266205

**Authors:** Mariann Oemisch, Stephanie Westendorff, Marzyeh Azimi, Seyed Ali Hassani, Salva Ardid, Paul Tiesinga, Thilo Womelsdorf

**Author notes:** Corresponding Authors: Dr. Mariann Oemisch, Dr. Thilo Womelsdorf.

## Abstract

Prediction errors signal unexpected outcomes indicating that expectations need to be adjusted. For adjusting expectations efficiently prediction errors need to be associated with the precise features that gave rise to the unexpected outcome. For many visual tasks this credit assignment proceeds in a multidimensional feature space that makes it ambiguous which object defining features are relevant. Here, we report of a potential solution by showing that neurons in all areas of the medial and lateral fronto-striatal networks encode prediction errors that are specific to separate features of attended multidimensional stimuli, with the most ubiquitous prediction error occurring for the reward relevant features. These feature specific prediction error signals (1) are different from a non-specific prediction error signal, (2) arise earliest in the anterior cingulate cortex and later in lateral prefrontal cortex, caudate and ventral striatum, and (3) contribute to feature-based stimulus selection after learning. These findings provide strong evidence for a widely-distributed feature-based eligibility trace that can be used to update synaptic weights for improved feature-based attention.

**Highlights:**

- Neural reward prediction errors carry information for updating feature-based attention in all areas of the fronto-striatal network.
- Feature specific neural prediction errors emerge earliest in anterior cingulate cortex and later in lateral prefrontal cortex.
- Ventral striatum neurons encode feature specific surprise strongest for the goal-relevant feature.
- Neurons encoding feature-specific prediction errors contribute to attentional selection after
learning.

## Introduction

When faced with novel objects we have to learn which features defining these objects are relevant, and which can be safely ignored. To succeed learning which visual features of objects are relevant likely depends on estimating feature relevance and improving this estimate through trial and error learning (Farashahi et al., 2017; Hikosaka et al., 2017; Wilson and Niv, 2011). Computational work shows that this improvement of estimated feature relevance can be achieved by calculating how unexpected an experienced outcome is, and updating its estimate in proportion to this unexpectedness (Leong et al., 2017; Niv et al., 2015). In typical reinforcement learning models, the unexpectedness is calculated as prediction error between predicted and experienced outcome value (Sutton and Barto, 1998; Watkins and Dayan, 1992).

For prediction errors to be useful they need to inform the subject about the specific feature causing the unexpected outcome. In fact, a prominent hypothesis suggests that the degree of unexpectedness and strength of prediction errors are guiding subject’s attention towards those features that gave rise to the unexpected outcome (Gottlieb, 2012; Pearce and Hall, 1980). Biasing attention to those features causing outcomes that have been most unexpected can optimize attentional sampling in the long run to those stimuli with the most reward-predictive features (Daddaoua et al., 2016; Dayan et al., 2000; Ghazizadeh et al., 2016). The mechanisms underlying this attentional optimization through reinforcement learning have been explored in recent studies suggesting that attentional guidance by prediction errors is facilitated when value predictions are already biased towards those feature dimensions that are most likely reward predictive (Hassani et al., 2017; Leong et al., 2017; Niv et al., 2015; Wilson and Niv, 2011). Instead of attending all possible feature dimensions of a stimulus equally, prioritizing those dimensions that most prominently are reward predictive dramatically enhances the learning speed (Farashahi et al., 2017; Kruschke and Hullinger, 2010). This learning speed increase is particularly prominent with stimuli that are defined by multiple dimensions as is typical of real world objects. A strong prediction of these behavioral models is that brain circuits need to combine information about the occurrence of a prediction error with information about the task relevant feature dimension that should be attended (Asaad et al., 2017). However, it is unknown how this combination of prediction error information and feature-based attention is realized in brain circuits.

Here, we set out to identify how this combination of prediction errors and task relevant stimulus features is encoded in medial and lateral fronto-striatal circuits composed of anterior cingulate and ventral striatum as the medial loop and the lateral PFC and caudate nucleus as the lateral loop (Haber and Knutson, 2010). In one scenario of learning which stimulus features are relevant, a general prediction error signal emerges locally within the ventral striatum and is broadcasted to prefrontal cortex where it modifies the activity of feature selective neurons(Asaad et al., 2017). Updated prefrontal cortex neurons might then exert an improved top-down signal over sensory cortices for attention and choices in subsequent trials (e.g. (Fusi et al., 2007; Seo et al., 2012). This view is supported by a ubiquity of mostly human fMRI findings that single out the striatum as core region to encode prediction errors (Chase et al., 2015; Glimcher, 2011), and the lateral prefrontal cortex to encode a feature-based top-down signal (Leong et al., 2017; Serences, 2008) together with prediction error information (Asaad et al., 2017). In contrast to such a view emphasizing functional localization, neurons encoding prediction errors in multiple areas might carry already feature information. Such feature specific prediction errors could serve as feature specific eligibility trace orchestrated across the recurrent fronto-striatal loops. Such a distributed, feature-specific eligibility trace is predicted by spiking network models that learn task relevant features by using attentional feedback signals to label synapses among those neurons that also contributed to the feature specific reward prediction itself (Roelfsema and van Ooyen, 2005; Rombouts et al., 2015).

Here, we tested these scenarios by recording from four brain areas of the medial and lateral fronto-striatal loops in macaque monkeys performing a feature-based value learning task. Task performance was well fit by an attention-weighting reinforcement learning model that estimated trial-by-trial prediction errors. We found that substantial proportions of neurons across all brain areas encoded prediction errors for task relevant features. The strongest encoding was evident for the task and reward relevant feature, which represents a versatile eligibility trace that can guide synaptic learning across the whole fronto-striatal loop. We found that this feature specific eligibility trace emerged after an initial unspecific prediction error signal and was associated with stronger attentional selection signals in subsequent trials, thus potentially contributing to improved learning and visual selection.

## Results

### Behavior

Monkeys performed a reversal learning task which presented in each trial two peripheral stimuli with different colors and motion directions (**Figure 1A**). Over sequences of 30 or more correct trials one of two colors was associated with reward outcomes (juice drops), while no other feature (left vs. right stimulus location, or up vs. downward motion direction, or the alternative color) was linked to reward (**Figure 1B**). In order to obtain reward the animals had to wait for a Go-signal (dimming of the stimuli) and make a saccade in the motion direction of the grating stimulus whose color matched the reward associated color. This task required (1) feature-based attentional selection of one over another stimulus based on a reward associated color, and (2) to use the motion direction of the attended stimulus to program a saccadic response. Above chance performance on this task required learning that a correct (rewarded) or incorrect (nonrewarded) outcome was caused by the attended color of the stimulus rather than by its motion direction or its location on the screen. Both monkeys learned this feature specific credit assignment and adjusted their feature based attention bias to the reward associated color after uncued reversals (**Figure 1C**). As estimated with an ideal observer statistic (Balcarras et al., 2016; Smith et al., 2004), monkey H / K successfully learned an average of 83 / 91% of blocks, whereby learning occurred on average within 17.5 / 16.5 number of trials following the reversal. Monkey H / K performed an average of 9.7±0.3 / 8.9±0.3 reversal blocks per recording session with an average block length of 46±0.7 (*monkey H*, median = 37) and 43±0.8 (*monkey K*, median = 36) trials.

**Figure 1.**
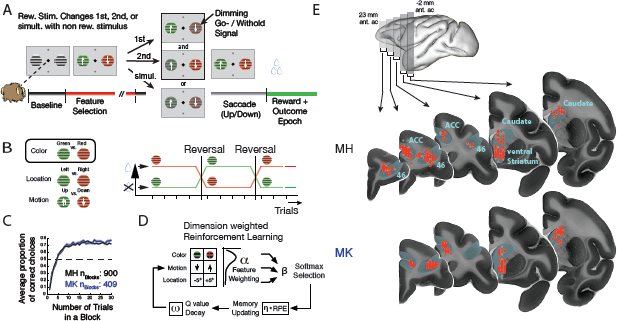
Feature-based reversal learning task and anatomical recording locations. (**A**) *Left:* Animals are presented with two black/white stimulus gratings to the left and right of a central fixation point. The stimulus gratings then become colored and start moving in opposite directions. Dimming of the stimuli served as Choice/Go signal. At the time of the dimming of the target stimulus the animals had to indicate the motion direction of the target stimulus by making a corresponding up- or downward saccade in order to receive a liquid reward. Dimming of the target stimulus occurred either before, after or at the same time as the dimming of the distractor stimulus. (**B**) *Left:* Three features characterize each stimulus – color, location, and motion direction. Only the color feature is directly linked to reward outcome. The task is a deterministic reversal learning task, whereby only one color at a time is rewarded. *Right:* This reward contingency switches repeatedly and unannounced in a block-design fashion. (**C**) Average proportion of correct choices relative to the reversal for monkey H (grey) and monkey K (blue). (**D**) Dimension weighted reinforcement learning model with main parameters α, β, ω, and η for feature weighting, selection noise, decay rate and learning rate, respectively. (**E**) Illustration of recording locations relative to stereotaxic zero for monkey H (top) and monkey K (bottom). Neuron locations are collapsed across 5mm coronal slices indicated by the grey bars on the brain on top. Red circles represent neurons that encoded a feature-specific prediction error, grey circles represent neurons that did not. Coronal images displayed are from an average macaque monkey (MRI atlas from (Calabrese et al., 2015).

### Neuronal Encoding of Outcome and Feature-Specific Reward Prediction Errors

During reversal learning performance, we recorded 1960 units in two monkeys with 690 units in the ACC (*monkey H:* 405, *monkey K:* 285), 524 units in PFC (*monkey H:* 316, *monkey K:* 208), 449 units in caudate nucleus (CD; *monkey H:* 234, *monkey K:* 215) and 297 units in ventral striatum (VS; *monkey H:* 163, *monkey K:* 134) (**Figure 1E**). 71% of neurons in *monkey H* and 78% of neurons in *monkey K* met the criteria for analysis (*see* Methods). Among these neurons 38% encoded the outcome (rewarded versus unrewarded), ranging between 27-53% across brain areas (**Suppl. Figure S1A, S3A, B**). To discern trial-by-trial encoding of prediction errors we fitted an attention weighting reinforcement learning model to the choice data of the monkeys (**Figure 1D**, *see* Methods) (Hassani et al., 2017; Leong et al., 2017; Wilson and Niv, 2011). We then correlated the model derived reward prediction errors (RPEs) with the neural firing rates during the 0-1.5 sec. reward outcome epoch. For monkeys H/K the firing rates correlated significantly with positive RPE (following correct trials) in 21/22% of neurons, with the negative RPE (following incorrect trials) in 14/10% of neurons, and for 24/24% of neurons with the unsigned RPE that indexes surprise (e.g. (Hayden et al., 2011)). Firing correlations with predictions errors were evident in each of the recorded brain areas in both monkeys (**Suppl. Figure S1B**).

We next hypothesized that in order to use prediction error information to adjust feature-based attention, neurons might encode RPEs not equally for all features of the stimulus, but selectively for the task relevant features. We found support for this suggestion in multiple example neurons with such feature selective encoding of RPE (**Figure 2**). For instance, the VS neuron in **Figure 2i** scales its firing rate with surprise (unsigned RPE) when color 1 was selected for the choice (top), but not when color 2 was selected (middle). The ACC neuron in **Figure 2iv** scales its firing rate with the negative RPE (greater firing with more negative RPE) when the selected stimulus was located on the left (top), but not when it was located on the right (middle). And the PFC neuron in **Figure 2v** scales its firing rate with the positive RPE, when the stimulus selected in the preceding choice was located on the right (top), but not when it was located on the left (middle). Overall, we found that 53.1% of neurons (*monkey H:* 52.7%, *monkey K:* 53.6%) across the fronto-striatal areas tested here encoded feature-specific positive, negative and unsigned surprise signals (**Suppl. Figures S6, S7**).

**Figure 2.**
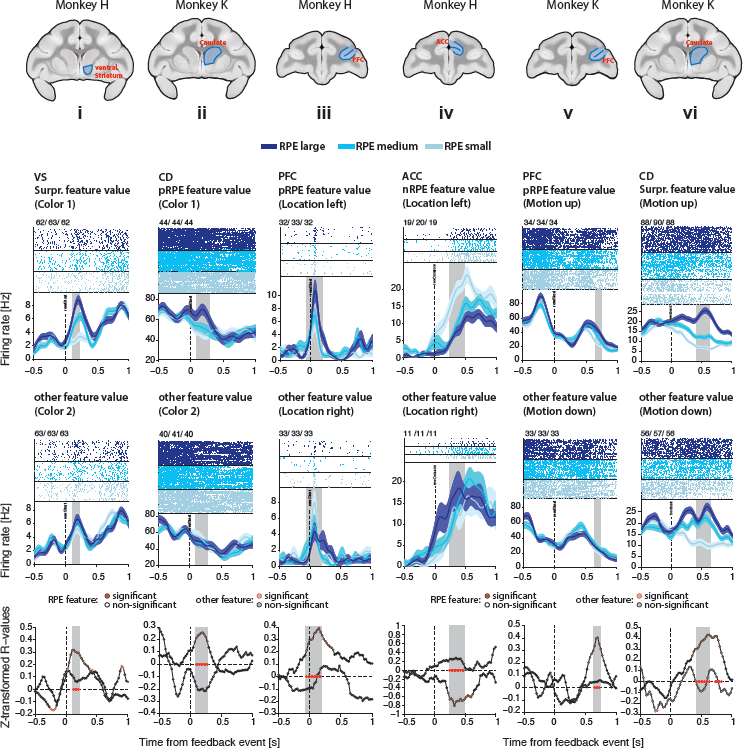
Example neurons encoding RPE signals for different feature- and RPE-types and from different areas and monkeys. For each of six example neurons (**i** - **vi**), the spike rasters and spike density functions are displayed (for visualization purposes only) for prediction error values of three different magnitudes (trials evenly split into RPE large, RPE medium, RPE small), for the feature value (e.g. color 1) for which an RPE was encoded (top row), and for the feature value for which an RPE was less or not encoded (e.g. color 2, middle row, ‘other feature value’). The bottom most row displays the z-transformed R values of the correlation between spike rate and RPE for the two feature values above (solely this last row displays the statistical analyses performed). Black empty (non-significant) or red filled (significant) circles represent z-transformed R values of the correlation between spike rate and RPE for those feature trials for which a RPE was encoded. Grey filled circles with black (nonsignificant) or red (significant) outlines represent z-transformed R values of the correlation between spike rate and RPE for those feature trials for which a RPE was *not* encoded. Red stars indicate those time bins for which the R-values between the two feature values differed significantly (Z-test, see Methods, eq. 5). Grey transparent bars in *all* plots indicate the time window of RPE encoding. The title above each column of figures indicates the area that neuron was recorded from as well as the type of feature and RPE signal encoded by that neuron. Anatomical images at the top-most additionally illustrate the recording locations. Shaded error bars represent SEM.

### Feature-specific RPE Signals Emerge Later than Non-specific RPEs and Earliest in ACC

Feature specific RPE signals might arise from neurons that initially encode the occurrence of a non-specific prediction error by combining RPE with feature information over time. This suggestion predicts a slower time course of more specific information about the source of the error (Schultz, 2016). We tested this possibility by determining for each neuron the time window in which it significantly encoded a feature-specific RPE, non-specific RPE, or outcome per se for four consecutive time bins (≥0.1 sec.), and comparing their average time-courses (see Methods). We found across neurons that feature specific RPE encoding emerged significantly slower than non-specific RPE encoding as indexed by a shallower slope of the temporal cumulation of the proportion of significantly RPE encoding neurons (Kolmogorov-Smirnoff test, Bonferroni-Holm multiple comparison corrected: p_feat-non_ < .001) (**Figure 3A, B**). Prediction errors in general emerged significantly later than the encoding of the rewarded/nonrewarded outcome (Kolmogorov-Smirnoff test, Bonferroni-Holm multiple comparison corrected: p_feat-non_ < .001; p_feat-out_ < .001; p_non-out_ < .001). In addition, feature specific RPE encoding continued to increase and remained at a higher plateau level for a longer duration than nonspecific RPE signals (**Figure 3A**). Across units, 25% of outcome encoding occurred at 268ms, while 25% of non-specific RPE encoding occurred 283ms after feedback onset, and 25% of feature-specific RPE encoding occurred after 355ms (Randomization statistic: p_feat-non_ < .001; p_feat-out_ < .001; p_non-out_ = .008; **Figure 3B**).

**Figure 3.**
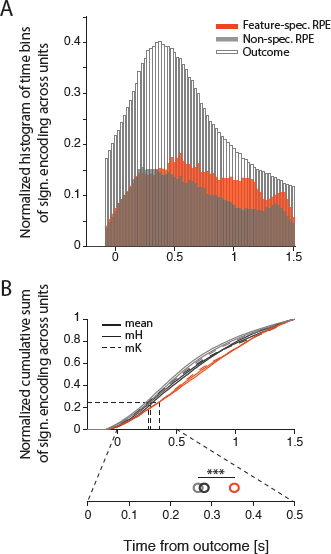
Temporal profile of feature-specific RPE, non-specific RPE and outcome signals. (**A**) Time courses of feature-specific RPE encoding, non-specific RPE encoding and outcome encoding across all units that encode the given signal combined across both monkeys. For each neuron, *all* time bins for which an RPE was encoded are included. (**B**) Normalized cumulative sums of the histograms in (A). *Top:* Thick lines represent the mean across both monkeys, while thin continuous lines represent cumulative sums of *monkey H*, and thin dotted lines represent cumulative sums of *monkey K*. All three cumulative sums differed significantly from each other (Kolmogorov-Smirnoff test, Bonferroni-Holm multiple-comparison correction; all p < .001). *Bottom:* Magnification of the cumulative sums around the 25% window. Open circles represent the time points at which 25% of the respective signal is encoded. The horizontal bar with three asterisks indicates that all three time points differ significantly from each other (randomization procedure, all p < .01).

We next ask when feature specific RPE encoding emerges in each of the four brain areas. Using the same latency measures as above, we found that the rise of neurons with significant feature specific RPE differed significantly between all areas, except for ACC and CD which did not differ (Kolmogorov-Smirnoff test, Bonferroni-Holm multiple comparison corrected: p_ACC-PFC_ < .001; p_ACC-CD_ = .128; p_ACC-VS_ < .001; p_PFC-CD_ < .001; p_PFC-VS_ = .006; p_CD-VS_ < .001) (**Figure 4A, B**). Feature-specific RPE signals emerged earliest in the ACC (310ms) and CD (330ms), followed by PFC (385ms), followed by VS (428ms) (Randomization statistic: p_ACC-PFC_ < .001; p_ACC-CD_ = .136; p_ACC-VS_ < .001; p_PFC-CD_ < .001; p_PFC-VS_ = .018; p_CD-VS_ < .001; **Figure 4B** bottom).

**Figure 4.**
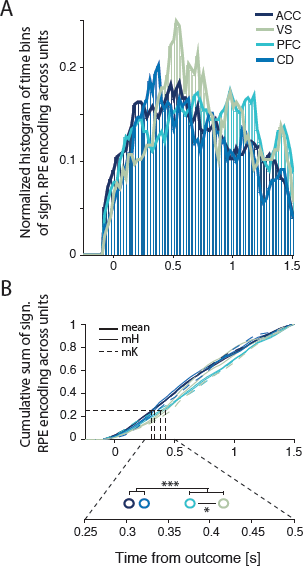
Latency comparison of feature-specific RPE encoding across areas. (**A**) Time courses of feature-specific RPE encoding in ACC, VS, PFC and CD neurons combined across both monkeys. To enhance visualization of the four histograms lines representing the outlines of each histogram are added. (**B**) Normalized cumulative sums of the histograms in (A). *Top:* Thick lines represent the mean across both monkeys, while thin continuous lines represent cumulative sums of *monkey H*, and thin dotted lines represent cumulative sums of *monkey K.* The cumulative sums of all areas except for ACC and CD differed significantly from each other (Kolmogorov-Smirnoff test, Bonferroni-Holm multiple-comparison correction; p_ACC-CD_ = .128, all other p < .01). *Bottom:* Magnification of the cumulative sums around the 25% window. Open circles represent the time points at which 25% of feature-specific RPEs is encoded in the four areas. One asterisk indicates p < .05; three asterisks indicate p < .001 (randomization procedure).

### Feature-tuning of Reward Prediction Errors

To guide reversal learning towards the goal-relevant color feature, subgroups of neurons should preferentially encode the prediction error for the reward-relevant color dimension. Such color specific error signals are likely candidates to update the task set representation following a reversal. In contrast to color, motion and location were choice-relevant stimulus dimensions that are important for completing the current trial selection, but do not facilitate reversal learning of the new color-reward rule. Consistent with this rationale, we found negative and positive RPEs were encoded more often for the reward-relevant color dimension, than for the location or motion dimension (one-sided bootstrap CI: p ≤ 0.05; **Figure 5A and D**, respectively). When split by areas, we found that neurons with color-specific negative RPEs were more prevalent than location- or motion-specific negative RPEs in ACC, VS, and PFC (one-sided bootstrap CI: p ≤ 0.05, **Figure 5B**). We used an index to quantify the relative proportion of color-selective RPE neuron compared to location- and motion-selective RPEs [(RPE_col_ – RPE_loc+motion_/2) / (RPE_col_ + RPE_loc_ + RPE_motion_)] with >0 values indicating a preference to encode RPEs for the goal relevant color feature. This color tuning index showed that for negative RPEs ACC, VS, and PFC showed stronger color tuned RPEs than CD (two-sided bootstrap CI: p ≤ 0.05; **Figure 5C**, see Methods). Similar to negative RPEs, positive RPEs were more often color-specific than location- or motion-specific in ACC and VS (one-sided bootstrap CI: p ≤ 0.05, **Figure 5E**, left column). In addition, neurons in CD also selectively encoded feature-specific positive RPEs in the color dimension, while neurons in PFC were not selective (**Figure 5E**, right column). Color tuning indices did not differ substantially between areas (ACC: I_col_ = 0.10, VS: I_col_ = 0.14, PFC: I_col_ = 0.05, CD: I_col_ = 0.123; two-sided bootstrap CI: p > 0.05; **Figure 5F**). The average correlation strengths between RPE and firing rate across time for those neurons that encoded a color-specific negative or positive RPE were comparable across areas (**Suppl. Figure S2**).

**Figure 5.**
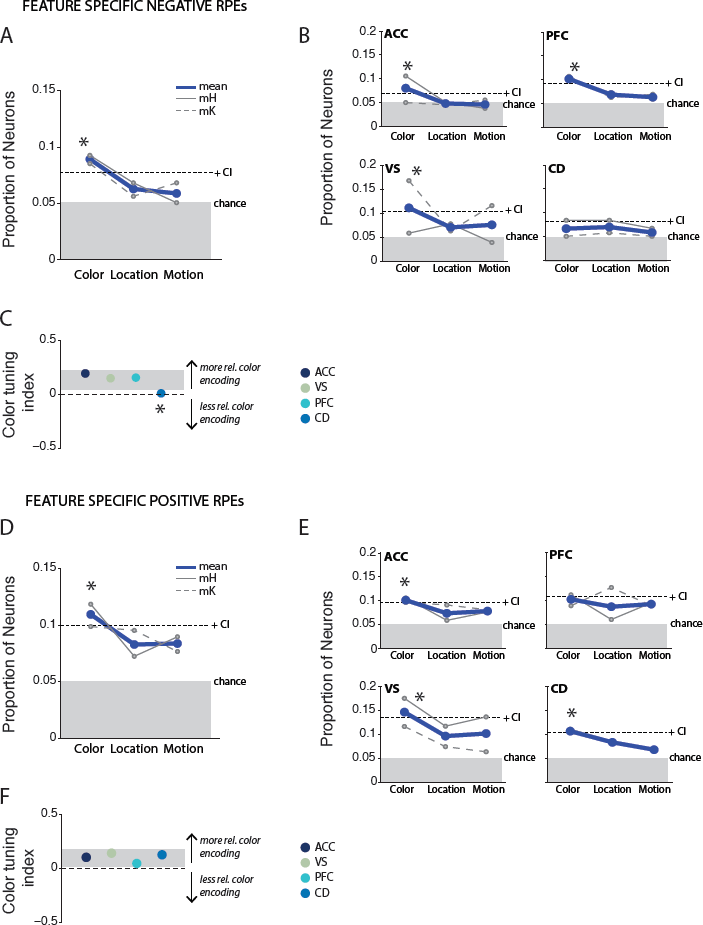
Prevalence of color-, location- and motion-specific negative and positive RPE encoding. Shown are proportions of neurons that encode a color-, location-, or motion-specific RPE negative signal either combined across areas (**A**) or split by areas (**B**). Thick blue lines represent averages across both monkeys. Thin continuous grey lines represent data from *monkey H*, thin dashed grey lines represent data from *monkey K.* An asterisk indicates p ≤ .05 using a one-sided bootstrap procedure that randomized the feature labels. Dotted lines indicate upper confidence interval. Grey bars indicate chance level proportion at 0.05. (**C**) Color tuning indices for each area computed according to eq. 6. Grey bar represents upper and lower bootstrap confidence interval. An asterisk indicates p < .05 by falling outside of the specified confidence interval. (**D-F**) equivalent conventions to (**A-C**) for feature-specific positive RPE encoding.

In contrast to positive and negative RPEs, unsigned surprise signals were across areas similarly prevalent for the color, location and motion dimensions (one-sided bootstrap CI: p > 0.05; **Figure 6A**). Split by areas, only the ventral striatum encoded surprise signals stronger for color than motion and location (one-sided bootstrap CI: p ≤ 0.05, **Figure 6B** bottom left). This finding was confirmed by a significantly higher color tuning index for VS (I_col_ = 0.103) than for ACC (I_col_ = -0.024), PFC (I_col_ = -0.036), and CD (I_col_ = -0.01) (**Figure 6C**). The average correlation strength of color-specific unsigned RPE units was similar across areas (**Suppl. Figure S2**).

**Figure 6.**
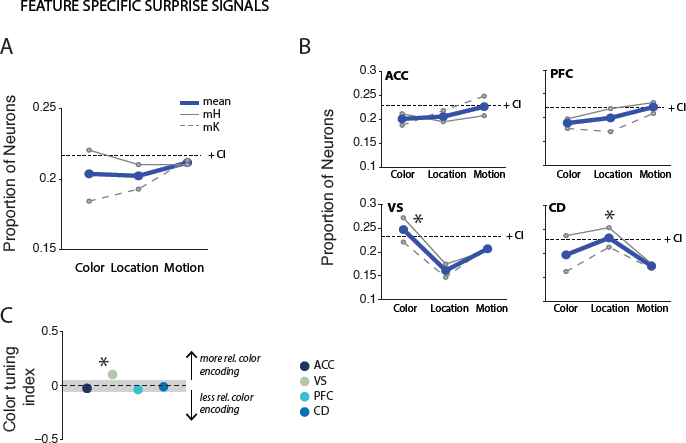
Prevalence of color-, location- and motion-specific surprise RPE encoding. Conventions are equivalent to Figure 7 for feature-specific unsigned RPEs.

### Feature-Specific RPEs are Largely Segregated from Feature-Specific Outcome Signals

Feature specific correlations of firing rates with the RPE signals might be evident in populations of neurons that show already feature specific firing for the outcome itself irrespective of prediction error, or they could occur in a segregated neuronal population. To answer this question, we first calculated the prevalence of feature information in the outcome period. We found that 20/16/13% of neurons encoded color/motion/location specific information in the outcome epoch at the first or second order (see **Methods** and **Suppl. Fig. S3**). However, only 35/28/26/% of these neurons also correlated their firing with color/motion/location specific RPEs, suggesting that a substantial population of feature-specific RPE encoding neurons cannot be explained by error independent feature preferences (**Suppl. Fig. S4**).

### Cell-type Specificity of RPE Encoding Neurons

To understand the mechanisms underlying feature specific prediction errors it will be important to identify the functional cell types encoding them. In our recordings, we used the action potential waveforms to distinguish two functional cell types in the cortical brain areas (putative pyramidal cells and putative interneurons), and two cell types in the striatum (putative medium spiny neurons and putative interneurons) using methods established before (Ardid et al., 2015; Berke, 2008; Lansink et al., 2010) (see **Methods and Fig 7**). In the cortical areas ACC and PFC, we found that narrow spiking neurons (putative interneurons) were more likely to encode feature-specific RPE signals (ratio narrow/broad = 0.65) than we would expect based on the population distribution of narrow to broad spiking units we recorded (ratio narrow/broad in population = 0.40) (Chi-square test, p ≤ .05), and that this was not the case for neurons encoding non-specific RPE signals (ratio narrow/broad = 0.53; Chi-square test, p > .05; **Fig 7C-E**). The same effect was visible as a statistical trend for the striatal areas, caudate and ventral striatum, whereby feature-specific RPEs were more frequently encoded by narrow spiking neurons (putative interneurons, (Berke et al., 2004; Kawaguchi, 1993; Lansink et al., 2010) than suggested based on the population distribution (Chi-square test feature-specific RPEs: p = .08) (**Fig 7H-J**).

**Figure 7.**
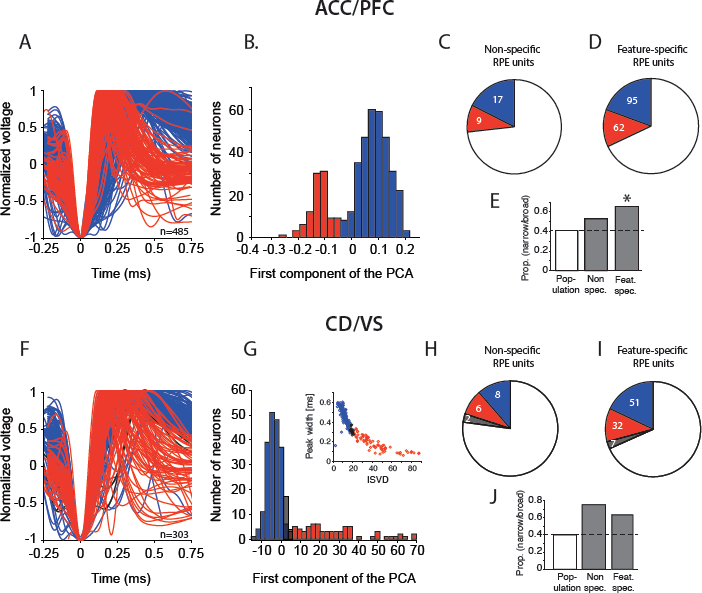
Cell-type classification of RPE units. (**A**)-(**E**) for ACC/PFC units. (**A**) Waveforms of all highly isolated single units recorded, identified as putative interneurons (narrow-spiking, red), putative pyramidal cells (broad-spiking, blue). (**B**) Histogram of the first component of the PCA using peak-to-trough duration and time to repolarization to separate neurons into putative interneurons and putative pyramidal cells. (**C**) Proportion of non-specific RPE encoding neurons identified as narrow- or broad-spiking. (**D**) Proportion of feature-specific RPE encoding neurons identified as narrow- or broad-spiking or unidentified. (**E**) Ratio of narrow to broad spiking neurons identified in the population, for nonspecific and feature-specific RPE encoding neurons. Black asterisk indicates p < 0.05 (chi-square test). (**F**)-(**J**) for CD/VS units. (**F**) Waveforms of all highly isolated single units recorded, identified as putative interneurons (red) or putative medium spiny neurons (MSNs, blue), or unidentified (black). (**G**) Histogram of the first component of the PCA using peak width and initial slope of valley decay (ISVD) to separate neurons into putative interneurons and MSNs. Inset shows the distribution of peak width to ISVD across neurons. (**H**) Proportion of non-specific RPE encoding neurons identified as putative interneurons or MSNs. (**I**) Proportion of feature-specific RPE encoding neurons identified as putative interneurons or MSNs or unidentified. (**J**) Ratio of putative interneuron/ MSN in the population, for non-specific and feature-specific RPE encoding neurons.

### Neuronal Signaling of Feature-Specific Prediction Errors can affect Stimulus Selection

What are the functional consequences of feature specific prediction errors? At the behavioral level, prediction errors indicate the need to adjust attention in subsequent trials. At the neuronal level, this adjustment for future attention might correspond to a shift of firing from the outcome time epoch early during learning to the firing at the time epoch of stimulus selection after learning. This temporal transfer of firing is the classical signature of reward prediction error encoding by ventral tegmental dopaminergic neurons (Fiorillo et al., 2003; Schultz, 1998; Schultz et al., 1993). To test whether such a transfer takes place for feature specific prediction errors, we determined whether the magnitude of the prediction error in the current trial was related to firing rate changes during stimulus selection in the following trial. We hypothesized that during learning periods when prediction errors are large, neurons would not yet contribute to the visual selection of the stimulus, but after learning when prediction errors are low, the same neurons would show an enhanced stimulus onset response indicating that they contribute to the attentional selection of the stimulus. For each color-specific RPE encoding neuron, we found the 25% of trials with the largest RPE and the 25% of trials with the smallest RPE, and compared the change in firing rate from pre- to post-color onset in those trials. On average, across these color-specific RPE encoding neurons we found an increased firing from pre- to post-color onset (t-test, p < .0001 for each RPE type). This increased color onset response was on average (1) stronger following trials with low RPE than high RPE and (**Figure 8A, B** lower versus upper histograms, and **Suppl. Fig S5**), and (2) stronger when the next trials choice was for the preferred color of the neurons (**Figure 8A, B** cyan versus grey histograms) The difference in firing rate change for low versus high RPE trials was statistically significant for neurons encoding positive RPE (paired t-test, p < .001; p_nonpref_ = .185) (**Figure 8A**), and for neurons encoding surprise (paired t-test, p < .001, p_nonpref_ = .065) (**Figure 8B**), but not for neurons encoding negative RPE (p = .089, p_nonpref_ = .291; data not shown).

**Figure 8.**
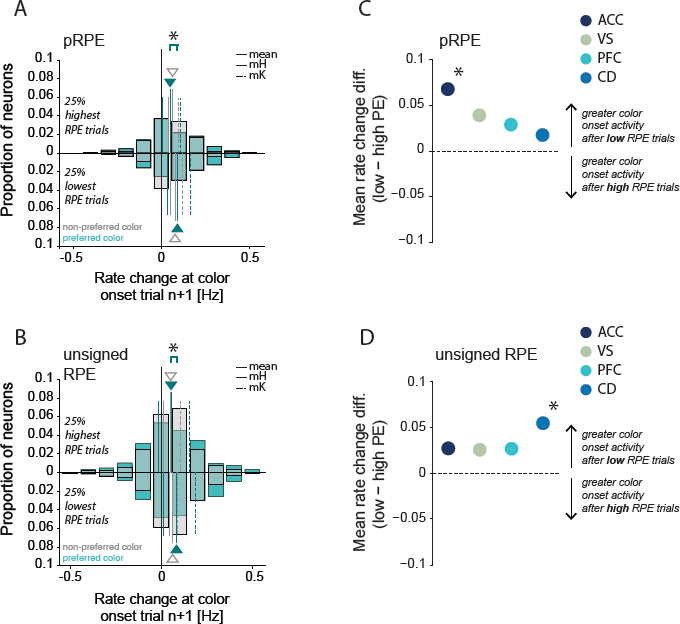
Firing rate increases in response to the color onset following low versus high RPE trials. (A,B) Firing rate changes following color onset in trial n+1 across populations of neurons encoding color-specific positive RPE (*A*) and surprise (unsigned RPE) (*B*). Above zero, rate changes are shown for the 25% of trials with the greatest prediction errors; below zero, rate changes are shown for the 25% of trials with the lowest prediction error, in cyan following preferred color choices and in grey following non-preferred color choices. Thick cyan and grey lines as well as triangles indicate average rate changes. Thin continuous lines show data from *monkey H*, thin dashed lines show data from *monkey K*. (C,D) Mean differences in rate changes following low versus high RPEs for each area areas across color-specific pRPE (*A*), and unsigned RPE (*B*) encoding neurons. Black asterisks indicate significant differences in rate changes following low versus high RPEs (paired t-test, p < .05).

The selectively increased firing to color onset after low RPE trials was most prominent and statistically significant for ACC neurons encoding color specific positive RPEs (**Fig 8C, Suppl. Fig S5**), and for CD neurons encoding color specific surprise (**Figure 8D, Suppl. Fig S5**). These finding provide strong evidence that feature specific prediction error signals during learning translate into color cue firing rate increases for these same neurons after learning has taken place, reminiscent of the temporal transfer of classical dopaminergic prediction error signals.

## Discussion

We found that about half of the neuronal populations in anterior cingulate cortex, lateral prefrontal cortex, ventral striatum and caudate nucleus encoded reward prediction errors specifically for one of three features of an attended visual stimulus during value learning. This feature specific prediction error was significantly stronger for the task relevant feature that also predicted reward over trials in a block (color), indicating that feature specific prediction error signals carry goal-relevant information that can bias improved feature-based attention in future trials. Feature specific prediction error encoding emerged in time after nonspecific prediction error encoding, indicating that it is based on a partly independent process that combines prediction error information with feature information over time.

### Prediction errors carry goal-relevant information to adjust feature-based attention

Among all recorded brain areas, the anterior cingulate cortex stood out by containing most neurons with early feature specific prediction error information, with a slower rise of feature information in the caudate head, lPFC, and ventral striatum (**Figure 4B**). This finding underlines the importance of the anterior cingulate cortex to indicate the specific information needed to adjust behavior in future trials (Shen et al., 2015; Shenhav et al., 2013; Shenhav et al., 2016). In our task, the goal-information was the specific color of the attended stimulus. This finding complements previous reports of anterior cingulate cortex neurons conveying prediction error related activity for specific actions (Matsumoto et al., 2007; Quilodran et al., 2008), unique objects (Kennerley et al., 2009), stimulus-response mapping rules (Johnston et al., 2007; Womelsdorf et al., 2010), attentional and motivational origin of errors (Shen et al., 2015), and more abstract combinations of stimulus and reward information (Kennerley et al., 2011). Our ACC finding uniquely adds to this literature by showing firstly, that ACC prediction error activity is combined with the attended color feature in an attention task that always presents all possible features on the screen, which induces perceptual ambiguity and uncertainty about stimulus features (**Figure 1**). Secondly, that this feature-specific prediction error activity potentially results in greater feature-based attention activity selectively for the preferred color with learning (**Figure 8**). These findings complement reports of attention specific activity in the ACC (Kaping et al., 2011; Oemisch et al., 2015; Voloh et al., 2015) supporting the view that ACC plays a major role for controlling to which of the available options (covert) attention shifts (Mesulam, 1981; Westendorff et al., 2016; Womelsdorf and Everling, 2015).

Thirdly, our findings clarify that specific information about the origin of prediction errors is not localized to the ACC, but widely distributes to all areas we recorded from and which are known to be anatomically mono-synaptically connected (Barbas and Pandya, 1989; Haber and Knutson, 2010; Hikosaka et al., 2017; Medalla and Barbas, 2009; Morecraft et al., 1993; Morecraft et al., 2012), and functionally synchronized in different task contexts (Antzoulatos and Miller, 2014; Oemisch et al., 2015; Voloh et al., 2015; Womelsdorf et al., 2014). The distributed nature of prediction error signaling supports recent summaries of the recurrent nature of fronto-striatal processes underlying reward-based choices (Hunt and Hayden, 2017), but also illustrates that latency analysis is able to identify a single brain area such as the ACC to have a particularly early functional contribution to value based attention and learning in a demanding reversal learning task used here.

The preponderance of narrow spiking neurons (in cortex) and as a trend of narrow spiking neurons in striatum to carry feature-specific error information provided an unexpected, data driven finding that support suggestions of a particular role of inhibitory neurons to process learning related information and/or to induce plasticity in cortical and striatal networks (Berke, 2011; Hennequin et al., 2017; Vogels et al., 2011). Previous studies suggest that the action potential waveform corresponds to inhibitory neurons in cortex and striatum (Kawaguchi, 1993; Plenz and Kitai, 1998; Wilson et al., 1994). Assuming that our finding indicate that putative interneurons are particularly informative about the error term is consistent with their involvement to regulate network level plasticity changes (Hennequin et al., 2017). Previous work has shown for instance that spike timing dependent plasticity in rodent corticostriatal excitatory synapses is crucially dependent on GABAergic signaling of inhibitory circuits (Paille et al., 2013), and that in balanced networks changes in inhibitory synaptic strength are accompanying changes in excitatory synaptic changes (Vogels et al., 2011).

### A role of surprise signals in the ventral striatum to guide ‘attention for learning’

For the ACC, PFC, CD and VS, prediction errors correlated with the firing of neurons after correct trials giving rise to positive RPEs, after incorrect trials giving rise to negative RPEs, and independent of the actual trial outcomes giving rise to unsigned unexpectedness, or surprise. Large surprise signals (to rare, high rewards) in the ACC have previously been shown to predict adjustment of behavioral strategies (Hayden et al., 2011), but it has been questioned whether any surprise related activity exists that relates to changes in attention (Le Pelley et al., 2016). Here, we found widely distributed and prevalent, neuronal surprise signals carrying significant information about all features of an attended stimulus to a similar degree in ACC, PFC, and CD. A notable exception was the ventral striatum which contained proportionally stronger neuronal surprise signals for the goal relevant color feature as opposed to the task relevant, but reward irrelevant, location and motion feature (**Figure 6C**). This preponderance of goal-relevant feature information in the ventral striatum is particularly noteworthy in light of a long-standing psychological theory of attention suggesting that attention during learning is driven by unexpected events (including outcome events) (Gottlieb, 2012, 2017; Pearce and Hall, 1980). According to this attention model, unexpected outcomes should give rise to stronger visual selection of the stimulus feature that gave rise to the violated expectation (Gottlieb et al., 2014). Our study directly tested this hypothesis and confirmed that the same neural population that encodes the feature specific surprise also showed stronger firing rate increases after the feature onset in subsequent trials (**Figure 8**). This increased feature selection signal (1) was stronger for the color that was preferred versus non-preferred by the neuron, and (2) it was stronger when subjects had learned the relevant feature, i.e. when prediction errors were comparably low in previous trials. These results provide strong evidence for a role of the ventral striatum to directly contribute to the learning of and attentional biasing towards goal relevant features. This conclusion adds an important functional role to prediction error signaling which is - across species - ubiquitously reported to be particularly strong in the ventral striatum (Chase et al., 2015; Cockburn et al., 2014; Costa et al., 2016; Diederen et al., 2016; Leong et al., 2017; O'Doherty et al., 2017; Schultz, 2017; Schultz et al., 2003; Takahashi et al., 2016; Watabe-Uchida et al., 2017).

Additionally, the finding of feature encoding in the ventral striatum and the caudate during learning of feature-based attentional allocation supports recent re-conceptualization of attentional biases as reflecting the internal striatal activity state that resolved competing value predictions and beliefs about possible relevant stimuli (Krauzlis et al., 2014; Womelsdorf and Everling, 2015). In these hypotheses, attention is not considered to reflect a unitary top-down signal that is obscure and localized to the prefrontal cortex as in many classical models, but rather attention emerges from (is the effect of) the current state of basal ganglia circuits that continuously integrate multiple information streams and resolves competition among these inputs (Krauzlis et al., 2014). A core insight from this hypothesis is that the striatum has direct access to the spatial maps in the superior colliculus via disinhibitory circuits in the substantia nigra (e.g. (Hikosaka et al., 2017)). With this direct access to the superior colliculus, activity in ventral striatum and caudate nucleus exert a direct bias for overt fixational sampling and covert attentional selection of visual information (Ignashchenkova et al., 2004; Lovejoy and Krauzlis, 2010; Zenon and Krauzlis, 2012). The results of our study support these hypotheses by revealing widespread feature specific prediction errors (**Figures 5** and **6**) and feature specific selection effects (**Figures 8** and **Supplementary Figure S5**) across the medial fronto-striatal loop (ACC and ventral striatum) and the lateral fronto-striatal loops (rostralateral PFC and dorsal caudate head).

### Neuronal credit assignment for relevant features with multidimensional stimuli

We report of neuronal populations encoding the ‘goal-specific’ reward prediction error for one color, but not for another color. These neuronal groups encode how unexpected it was that the attended color led to a reward, or to omission for reward. This information is precisely what is needed to enhance those synaptic connections between neurons that encode the specific color that is more relevant than expected, and to reduce the synaptic connection weights among neurons encoding the color that was less rewarded than expected. These types of synaptic weight changes following the strength of prediction errors have been successfully modelled in several spiking network models implementing reinforcement learning using different synaptic learning rules (Fremaux et al., 2013; Friedrich et al., 2011; Fusi et al., 2007; Potjans et al., 2011; Rasmussen et al., 2017; Rombouts et al., 2015; Santoro et al., 2016; Seung, 2003; Soltani and Wang, 2010; Suri and Schultz, 1998; Urbanczik and Senn, 2009). These models illustrate, for example, that simpler stimulus-response reversal learning performance in monkeys can be realized by spike timing dependent plasticity changes (Fusi et al., 2007). However, it has remained unclear how to implement more complex credit assignment in a higher dimensional feature space where multiple features could be credited for an outcome, even though only one feature is actually relevant (Niv et al., 2015; Wilson and Niv, 2011). For this situation, a recent spiking model suggested a 4-factor learning rule that is dependent on attention to a specific stimulus feature or action prior to registering a reward/no-reward outcome (Rombouts et al., 2015). In this model, neurons activated by an outcome receive a synaptic tag, which is specific to the attended feature, from feedback connections originating from output neurons similar to striatal output neurons. This attentional feedback induced synaptic tag acts like an attention specific eligibility trace that can be combined with dopamine dependent (feature unspecific) prediction error information when a (rewarding or non-rewarding) outcome is received. Learning is achieved when these two factors (attentional feedback and neuromodulatory prediction error information) meet at the synapses between pairs of neurons that showed near coincident pre- and postsynaptic activity during the outcome processing (Rombouts et al., 2015). The models make multiple predictions that are supported by our data. Firstly, synaptic updating is taking place in an associative network layer that resembles the fronto-striatal network of value learning as opposed to sensory or motor related network layers. Secondly, feature specific prediction errors are predicted to emerge as local neuronal signals across the entire associative network based on neuron-specific synaptic tags, closely corresponding to the distributed RPEs we observed. Finally, the model predicts that a learning of task relevant features depends on attention towards those stimulus features that are most consistently reward associated. This attentional hypothesis of reinforcement learning was directly tested in our experiment, providing evidence that the most ubiquitously encoded prediction error signals occur for the attended, goal-relevant color feature.

Taking together, we believe that our findings provide direct support of the concept of attention weighted reinforcement learning as a generic framework to understand learning and attention in environments with multidimensional stimuli, as is typical for real life learning of object relevance (Hassani et al., 2017; Leong et al., 2017; Niv et al., 2015). The existence of network-wide available information about the degree to which individual features of visual stimuli led to unexpected outcomes will further constrain the modeling of learning rules that efficiently solve the credit assignment problem (Farashahi et al., 2017; Hennequin et al., 2017; Watabe-Uchida et al., 2017). Our study may provide a starting point documenting that network wide credit assignment processes are directly related to improved biases of feature-based visual attention. It will be an important question for future research to specify how feature specific eligibility traces are used to determine the value of objects during fast and slow learning.

## Acknowledgement

This work was supported by grant MOP 102482 from the Canadian Institutes of Health Research (TW) and by the Natural Sciences and Engineering Research Council of Canada (TW), and the Brain in Action CREATE-IRTG program (MO, TW). The funders had no role in study design, data collection and analysis, the decision to publish, or the preparation of this manuscript. Authors would like to thank Hongying Wang, Samira Azimi and Ali Hassani for technical support.

## Author contributions

Conceptualization, T.W.; Methodology, T.W., S.W. and M.O.; Formal Analysis, M.O., T.W., S.A. and P.T; Investigation, M.O., M.A., S.A.H.; Writing – Original draft, M.O. and T.W.; Writing – Review & Editing, all authors; Supervision, T.W.; Funding acquisition, T.W.

## Declaration of interests

The authors declare no competing interests.

## Methods

### Electrophysiological recordings

Data was collected from two male rhesus macaques (*Macaca mulatta*). All animal care and experimental protocols were approved by the York University Council on Animal Care and were in accordance with the Canadian Council on Animal Care guidelines. Extra-cellular recordings were made with 1-12 tungsten electrodes (impedance 1.2 - 2.2 MOhm, FHC, Bowdoinham, ME) in anterior cingulate cortex (area 24, ACC), prefrontal cortex (area 46, PFC), caudate nucleus (CD) and ventral striatum (VS) through rectangular recording chambers (20 by 25 mm) implanted over the right hemisphere (see **Figure 1E** and Suppl. Methods for anatomical reconstruction). Electrodes were lowered daily through guide tubes using software controlled precision microdrives (NAN Instruments Ltd., Israel). Data amplification, filtering, and acquisition were done with a multi-channel acquisition processor (*Neuralynx).* Spiking activity was obtained following a 300 - 8,000 Hz passband filter and further amplification and digitization at 40 kHz sampling rate. Sorting and isolation of single unit activity was performed offline with Plexon Offline Sorter, based on principal component analysis of the spike waveforms. Experiments were performed in a custom-made sound attenuating isolation chamber. Monkeys sat in a custom-made primate chair viewing visual stimuli on a computer monitor (60Hz refresh rate, distance of 58cm). Eye positions were monitored using a video-based eye-tracking system (*EyeLink, SRS Systems*) calibrated prior to each experiment to a 9-point fixation pattern. Eye fixation was controlled within a 1.4-2.0 degree radius window. During the experiments, stimulus presentation, monitored eye positions, and reward delivery were controlled via MonkeyLogic (http://www.brown.edu/Research/monkeylogic/). Liquid reward was delivered by a custom-made, air-compression controlled, mechanical valve system.

### Anatomical reconstruction

Recording locations were identified using MRI images obtained following initial chamber placement. During MR scanning, we placed a grid marking the chamber center and peripheral positions as well as a diluted iodine solution inside the chamber for visualization. This allowed the referencing of target regions to the chamber center in the resulting MRI images. Target regions (area 24 – ACC, area 46 – PFC, caudate nucleus, ventral striatum) were identified using the scheme from the Price lab (Saleem et al., 2008; Saleem et al., 2014). The dorsal-ventral positioning of electrodes was estimated daily using the MRI images and audible profiles of spiking activity. The relative coarseness of the MRI images did not allow us to differentiate recording locations in the shell of the nucleus accumbens as opposed to the core of the nucleus accumbens with certainty.

### Behavioral paradigm

The monkeys performed a feature-based reversal learning task that required covert spatial attention to one of two stimuli dependent on color-reward associations (**Figure 1A**). These color-reward associations were reversed in an uncued manner between blocks of trials with constant color-reward association (**Figure 1B**). By separating the location of attention from the location of the saccadic response, this task allowed an identification of neural responses to the location of attentional focus independent of neural signals linked to response preparations, during reversal learning. Each trial started with the appearance of a grey central fixation point, which the monkey had to fixate. After 0.5 - 0.9s, two black/white drifting gratings appeared to the left and right of the central fixation point. Following another 0.4s the two stimulus gratings either changed color to black/green and black/red (*monkey K*: black/cyan and black/yellow), or started moving in opposite directions up and down, followed after 0.5 - 0.9s by the onset of the second stimulus feature that had not been presented so far, e.g. if after 0.4s the grating stimuli changed color then after another 0.5 - 0.9s they started moving in opposite directions. After 0.4 - 1s either the red and green stimulus dimmed simultaneously for 0.3s or they dimmed separated by 0.55s, whereby either the red or green stimulus could dim first. The dimming represented the go-cue to make a saccade to one of two response targets displayed above and below the central fixation point. Please note that the monkeys needed to keep central fixation until this dimming event occurred. A saccadic response following the dimming was only rewarded if it was made to the response target that corresponded to the movement direction of the stimulus with the color that was associated with reward in the current block of trials, e.g. if the red stimulus was the currently rewarded target and was moving upward, a saccade had to be made to the upper response target at the time the red stimulus dimmed. A saccadic response was not rewarded if it was made to the response target that corresponded to the movement direction of the stimulus with the non-reward associated color. A correct response was followed by 0.33ml of water delivered to the monkey’s mouth. Across trials of a block the color-reward association remained constant for 30 to a maximum of 100 trials. Performance of 90% rewarded trials (calculated as running average over the last 12 trials) automatically induced a block change. The block change was un-cued, requiring the subject to use the reward outcome they received to learn when the color-reward association was reversed in order to covertly select the stimulus with the rewarded color. In contrast to color, other stimulus features (motion direction or stimulus location) were only randomly related to reward outcome. Saccadic responses had to be initialized within 0.5 s after dimming onset to be considered a choice (rewarded or non-rewarded). All other saccadic responses, e.g. towards the peripheral stimuli, were considered non-choice errors.

We used blocksine gratings with rounded-off edges for the peripheral stimuli, moving within a circular aperture at 0.8 °/s and a spatial frequency of 1.2 (cycles/°) and a radius of 2.0°. Gratings were presented at 5° eccentricity to the left and right of the fixation point.

### Data analysis

Analysis was performed with custom MATLAB code (Mathworks, Natick, MA), utilizing functions from the open-source Fieldtrip toolbox (http://www.ru.nl/fcdonders/fieldtrip/). All spike-density functions were smoothed with a Gaussian kernel with a standard deviation of 25ms. Only correct and incorrect choice trials were analyzed, whereby correct choice trials were rewarded trials, while incorrect choice trials were either made to the non-rewarded stimulus or in the incorrect response time window (first vs. second dimming). Fixation breaks, early responses, and no-response trials were not included in any analyses.

#### Initial neuron selection criteria

Units were only included in any of the following analyses if they i) had a minimum firing rate of 0.5Hz within the feedback epoch (0 - 1.5seconds following feedback onset), ii) prediction errors computed with a reinforcement learning model (see below) could be computed for ≥40 trials, and iii) these minimum of 40 trials could be identified as either occurring *during* learning or *after* learning according to an ideal observer statistics (see below). All trials from blocks that were not learned to criterion were discarded.

#### Expectation maximization algorithm

To identify at which trial during a block the monkey showed statistically reliable learning we analyzed the monkeys’ trial-by-trial choice dynamics using the state–space framework introduced by (Smith and Brown, 2003), and implemented by (Smith et al., 2004). This framework entails a state equation that describes the internal learning process as a hidden Markov or latent process and is updated with each trial. The learning state process estimates the probability of a correct (rewarded) choice in each trial and thus provides the learning curve of subjects. The algorithm estimates learning from the perspective of an ideal observer that takes into account all trial outcomes of subjects’ choices in a block of trials to estimate the probability that the outcome in a single trial is correct or incorrect. This probability is then used to calculate the confidence range of observing a correct response. We defined the learning trial as the earliest trial in a block at which the lower confidence bound of the probability for a correct response exceeded the p = 0.5 chance level. The identification of a learning trial allowed to discard blocks that were not learned.

#### Quantifying prediction errors with reinforcement learning modeling

We quantified the trial-by-trial progression of RPEs during reversal performance using a computational model that combines reinforcement learning (RL) principles with Bayesian tracking of reward probabilities for target features. This hybrid Bayesian-RL model was introduced before (Wilson and Niv, 2012; Niv et al., 2015) to account for behavioral adjustments of choices in a multidimensional visual learning task and was recently validated as a model accounting for feature-based reversal learning in the macaque (Hassani et al., 2017). The model represents the stimuli in terms of their stimulus features (color, motion, location), feature values (color A, color B, downward motion, upward motion, left, right), and the actual combinations of feature values for stimulus 1 and stimulus 2.

The model uses Bayesian inference about which stimulus feature f (color, motion or location) is the likely target feature via 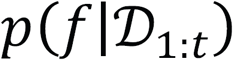 to obtain a feature-weighted representation for each stimulus. For tracking target feature probability, we denote the feature dimension as f_d_ (1: location, 2: direction of motion, 3: color) and for each d, f_d_, takes two values 1 and 2. For instance, f3=1 indicates the first color. We can then calculate the probability for the rewarded stimulus (the target) to have feature d, 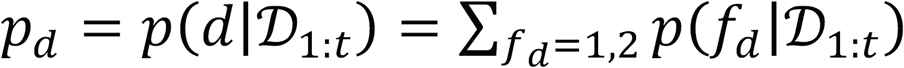. This defines a feature dimension weight 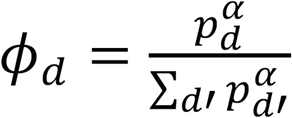 with exponent a and normalized to yield a sum across dimensions equal to one. The predicted reward value of a feature value is then denoted by 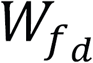 and scaled by the dimensional weight *ϕ_d_*. The value of the specific stimulus i is given by the sum across all weighted feature values that are part of the stimulus

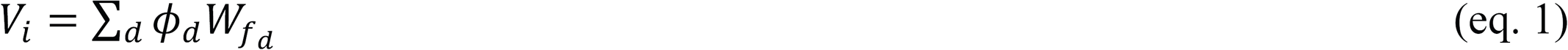

The choice of which stimulus is selected on a given trial is implemented with a softmax rule using the Boltzmann function

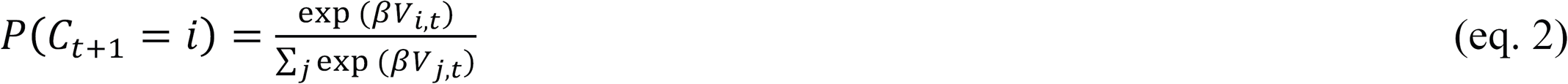

Following a choice, the stimulus values of the chosen stimulus are updated by a reward prediction error scaled by learning rate *η* according to:

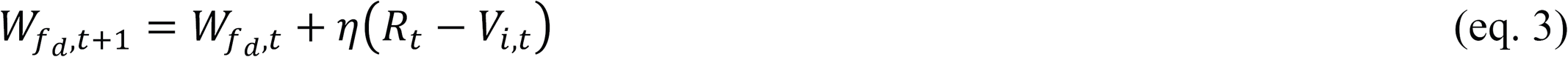

Values of the unchosen stimulus feature values were scaled down (decayed) by (1 – *ω*), similar to previous studies (see Hassani et al., 2017; Niv et al., 2015), according to:

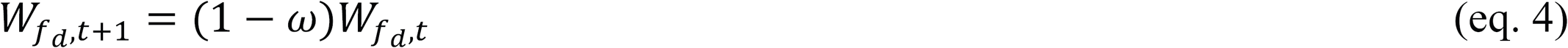

In summary, feature values of the chosen stimulus are updated using the RPE (eq. 3) and are separately scaled by a dimensional weight (that may be called attentional weight) calculated using Bayes updating of how the feature dimensions color, motion and location relate to reward outcomes.

We optimized the model by minimizing the negative log likelihood over all trials using up to 20 iterations of the simplex optimization method (matlab function fminsearch) followed by fminunc which constructs derivative information. We used an 80% / 20% (training dataset / test dataset) cross-validation procedure repeated for n=100 times to quantify how well the model predicted the data. Each of the one hundred cross-validations optimized the model parameters on the training dataset. We then quantified the log-likelihood of the independent test dataset given the training datasets optimal parameter values. We validated that the described hybrid Bayes-RL model provides a better fit (lower log-likelihood and lowest Akaike Information Criterion) for the cross-validated test dataset than simpler models that either lacked the Bayesian dimension weighting, or that lacked the decay of nonchosen stimulus features (for a detailed evaluation of different models, see also (Hassani et al., 2017)).

Both monkeys choice data were fit well by the Bayes-RL with log likelihoods for *monkey H* and *monkey K* of 0.47 and 0.52, respectively). The model parameters best explaining the data for *monkey H/K* had a similar pattern with η (learning rate) = 0.22/0.25, β (selection noise) = 3.55/2.79, ϕ (dimension weighting of feature representation) = 0.68/0.98 and ω (value decay for nonchosen feature) = 0.92/0.68. These results resonate well with previous studies using a similar model architecture (Wilson and Niv, 2012; Niv et al., 2015; Hassani et al., 2017; Leong et al., 2017).

#### Identification of prediction error encoding neurons

To identify RPE encoding neurons, we correlated each neuron’s firing rate time-resolved with RPEs obtained from the RL model. Each correlation analysis required a minimum of 15 trials. We correlated firing rate with positive RPEs in correct choice trials and with negative RPEs in incorrect choice trials. To identify neurons that encoded an unsigned RPE, we used partial correlation analysis to correlate firing rates with the absolute RPE in correct and incorrect choice trials while partializing out the sign of the RPE (by including a co-variate of +/-1 for correct/incorrect trials respectively). The analysis time ranged from -500 to 1500ms after the outcome event; time windows spanned 200ms and were shifted by 25ms. For a neuron to be considered to encode a non-specific positive, negative, or unsigned RPE signal, it had to significantly positively correlate its firing rate with a positive, or unsigned RPE, respectively (Spearman correlation, p < 0.05), for a minimum of four consecutive time bins following the outcome event, while not correlating positively in more than two consecutive time bins before the outcome event. For a neuron to be considered to encode a negative RPE signal, it had to significantly negatively correlate its firing rate with a negative RPE for a minimum of four consecutive time bins following the outcome event (Spearman correlation, p < 0.05), while not correlating negatively in more than two consecutive time bins before the outcome event.

To identify neurons that encoded a feature-specific RPE signal, trials were split into the features of interest prior to the correlation analysis (color, location and motion direction). The principle for identifying positive, negative and unsigned feature-specific RPE neurons was the same as for non-specific RPE signals with additional criteria described in the following. For instance, for a neuron to be considered to encode a color-specific RPE signal, it had to significantly encode a RPE signal (as described above) in minimally four consecutive time bins for trials in which e.g. color 1 was chosen, while either not encoding or encoding significantly less a RPE signal for trials in which color 2 was chosen. Significant differences between R values (Spearman correlation) for the two trial types were computed by z-transforming R values and comparing them using a z-test:

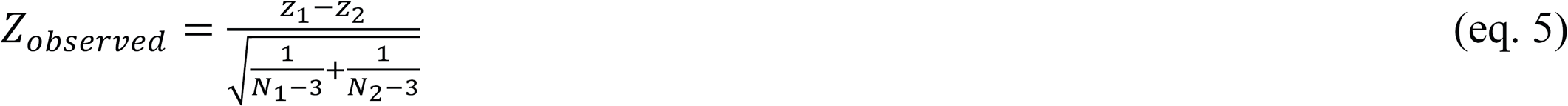

where z_1_ and z_2_ are the z-transformed R-values for the correlation with feature value 1 and feature value 2, respectively. When Z_observed_ exceeded |1.96| (p<0.05), R values were considered significantly different for a given time bin. In a minimum of four consecutive bins, R values from correlations with two different feature values (e.g. color 1 chosen or color 2 chosen) had to significantly differ, while a RPE had to be encoded for at least one of the two feature values according to the same criteria as for non-specific RPE signals. The method of identification was the same for identifying location and motion-specific RPE signals, with the exception of splitting trials according to chosen location or chosen motion direction, respectively. We determined for each neuron the duration in which it encoded a RPE signal as the first span of four or more consecutive significant time bins after the feedback event.

#### Time courses of prediction error and trial outcome signal encoding

To compare time courses of RPE signals, as well as trial outcome signals, we determined for each neuron the time window (minimum 4 consecutive bins) in which it encoded a RPE/trial outcome signal significantly (if a neuron encoded an RPE/trial outcome signal over longer time spans with time bins in between that were not significant, only the first time window of consecutive significant time bins was considered for this analysis). Across neurons, we therefore obtained distributions of time bins in which RPE/trial outcome signals were encoded, and we then tested these distributions for differences in their cumulative sums (Kolmogorov-Smirnoff test, Bonferroni-Holm multiple comparison correction, *α* = 0.05). As an additional measure of latency, we tested whether the time point at which 25% of RPE/trial outcome signals were encoded (the time point when the respective cumulative sum reaches 25%) differed using a randomization procedure (a = 0.05). The analysis procedure was equivalent when comparing the latencies of feature-specific RPE encoding between areas.

#### Comparing the prevalence of prediction error encoding

We used a bootstrap procedure to determine whether any feature-specific RPEs (color, location, or motion direction) were encoded more prevalently than would be expected based on the distribution across all feature-specific RPEs independent of their specificity (n=10,000). This bootstrap procedure was computed across all units encoding a specifically signed or unsigned RPE, initially independent of area recorded, and in a second step separately for each area. Significance was determined based on the actual proportion of color-, location-, or motion-specific RPEs falling inside or outside of the one-sided confidence interval of the bootstrap procedure. To compare the ratio of color-specific RPE encoding versus location- or motion-specific RPE encoding between areas, we computed a color tuning index for each area as follows:

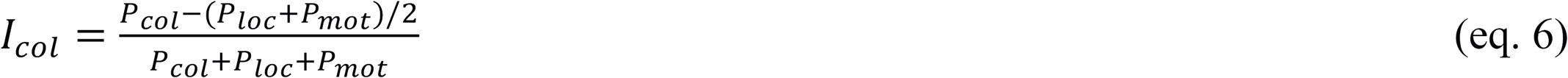

whereby I_col_ refers to the color tuning index, P_col_, P_loc_, and P_mot_ refer to the proportions of color-, location-, and motion-specific RPE units, respectively. We then compared color tuning indices across areas by computing a confidence interval (bootstrap procedure, n=10,000) around color-tuning indices that were computed with randomized area labels. An area was considered to have a significantly greater or smaller color tuning index than the other areas if it fell outside of the confidence interval.

#### Cell-type specificity of RPE encoding neurons

For the set of highly isolated neurons (*monkey H:* n = 428, *monkey K:* 398), we aligned, normalized, and averaged all action potentials (Ardid et al., 2015). To distinguish putative interneurons (narrow-spiking) and putative pyramidal cells (broad-spiking) in PFC and ACC, we analyzed the peak-to-trough duration and the time for repolarization for each neuron (Oemisch et al., 2015). The time for repolarization was defined as the time at which the waveform amplitude decayed 15% from its peak value. We computed the principal component analysis (PCA) and used the first component because it allowed for better discrimination between narrow- and broad-spiking cells, compared to any of the two measures alone (Hartigan dip test, p < 0.0005). In addition, a comparison of Akaike’s and Bayesian was used to confirm that a two-Gaussian model fit the data better than a one-Gaussian model. To distinguish putative interneurons and putative medium-spiny neurons (MSNs) in CD and VS, we analyzed the peak width (at half maximum) and *I*nitial *S*lope of *V*alley Decay (ISVD, Berke, 2008; Lansink et al., 2010), as they provided a better waveform discrimination than e.g. peak-to-trough duration. The ISVD was computed as follows:

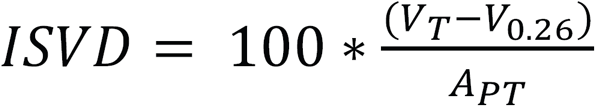

where V_T_ is the most negative value (trough) of the spike waveform, V_0.26_ is the voltage at 0.26 ms after V_T_, and A_PT_ is the peak-to-trough amplitude (Lansink et al., 2010). Although we could not discard unimodality for the first PCA component (or for either of the single measures, Hartigan dip test, p > 0.05), Akaike’s and Bayesian information criteria confirmed that a two-Gaussian model fit the data better than a one-Gaussian model. For frontal and cingulate units, we then used the two-Gaussian model and divided neurons into two groups of narrow and broad spiking units. For striatal units, because we could not discard unimodality for the first PCA component, we used the two-Gaussian model and defined two cutoffs that divided neurons into three groups. The first cutoff was defined as the point at which the likelihood of a narrow-spiking/putative interneuron was 3 times larger than the likelihood of a broad-spiking/putative principal cell, and vice versa for the second cutoff. We reliably classified PFC/ACC neurons (n = 485) as either putative pyramidal cells (broad spiking, n = 344, *monkey H:* 203, *monkey K:* 183) or putative interneurons (narrow-spiking, n = 141, *monkey H:* 78, *monkey K:* 49). Therefore, in *monkey H* 72% of neurons in ACC/PFC were identified as putative pyramidal cells while 28% of neurons were identified as putative interneurons. In *monkey K*, 82% of neurons were identified as putative pyramidal cells and 17% as putative interneurons. We classified 96% of striatal neurons (n = 277) as either putative MSNs (broad spiking, n = 198, *monkey H:* 96, *monkey K:* 113) or putative interneurons (narrow-spiking, n = 79, *monkey H:* 35, *monkey K:* 36), while n = 26 (*monkey H:* 8, *monkey K:* 11) neurons fell in between the criteria and could not be reliably classified. Therefore, in *monkey H* 73% of neurons in CD/VS were identified as putative MSNs while 27% of neurons were identified as putative interneurons. In *monkey K*, 77% of neurons were identified as putative MSNs and 23% as putative interneurons. For striatal units, we additionally verified our classification by comparing the firing rates between neurons classified as MSNs and those classified as interneurons. Striatal interneurons tend to be fast-spiking interneurons and should have a higher firing rate than the relatively low-firing MSNs (Berke et al., 2004; Berke, 2008). Indeed, in both monkeys, neurons classified as interneurons had a higher mean firing rate (*monkey H:* 4.96±1.1 Hz, *monkey K:* 4.77 ± 2.62 Hz) than neurons classified as MSNs (*monkey H:* 1.70 ± 0.38 Hz, *monkey K:* 1.61 ± 0.26 Hz), and this was statistically reliable in both monkeys (t-test, *monkey H:* p < 0.001, *monkey K:* p = 0.039). For the analysis of narrow versus broad spiking feature-specific versus non-specific RPE units we combined data from both monkeys because of relatively low neuron numbers. Proportions of narrow versus broad spiking units between non-specific and feature-specific RPE neurons were compared using chi-square statistics.

#### Stimulus selection following low and high prediction errors

We tested how neurons that encoded a color-specific prediction error changed their firing rate during color selection in trials following low versus high prediction errors (**Figure 7**). To do so, we identified for each color-specific RPE neuron the 25% of trials with the greatest prediction errors (from the model) and those 25% with the lowest prediction errors. These trials were then split into whether the choice was made to the color for which an RPE was encoded (preferred color) and those for which a choice was made to the other color (non-preferred color). For each trial n we found trial n+1 and computed the change in firing rate from the 400ms prior to stimulus color onset to the 100-700ms following stimulus color onset (post-color – pre-color / post-color + pre-color). We thus computed for each neuron the average rate change following low RPE trials in which the preferred color was chosen, following low RPE trials in which the non-preferred color was chosen, following high RPE trials in which the preferred color was chosen, and following high RPE trials in which the non-preferred color was chosen. Across the populations of color-specific positive RPE, negative RPE, or unsigned RPE encoding neurons, we then compared the change in firing rate at stimulus selection following high versus low RPE trials for the preferred color and for the non-preferred color choice trials using paired t-tests. In a second step, we split neuron populations based on their respective recording locations and performed the equivalent analysis.

#### Task variables encoded in the outcome epoch

To characterize neural responses in the outcome epoch, we adapted analyses from Padoa-Schioppa and colleagues (Cai and Padoa-Schioppa, 2014; Padoa-Schioppa and Assad, 2006, 2008). We tested whether neurons encoded any of twelve variables at the time of reward onset/omission. These twelve variables included the three stimulus features (color, location, motion) i) selected in the current choice independent of choice outcome (correct and error) (**chosen color, chosen location, chosen motion**) (Genovesio et al., 2014), ii) selected in the previous choice (trial n-1) independent of choice outcome (correct and error) (**previous chosen color, previous chosen location, previous chosen motion**) (Donahue and Lee, 2015; Genovesio et al., 2014), iii) of the target independent of choice (correct and error) (**target color, target location, target motion**) (Westendorff et al., 2016), in addition to the variables **outcome** (correct and error), **previous outcome** (correct and error) (Donahue and Lee, 2015) and **learning progress** (correct trials *during* learning versus *after* learning as obtained from the EM algorithm described above). To estimate the correlation between variables, we computed the correlation coefficient between any two trial vectors of the variables per recording session and then computed the average absolute correlation coefficient across recording sessions (the average correlation coefficient now varies between 0 and 1). The correlation matrix is shown in **Suppl. Fig. S3C**. To identify whether any neuron encodes any one variable, we performed independent linear regressions for each neuron on each variable. A neuron’s firing rate was averaged in the 0.1 - 0.7 seconds after reward onset/omission and was considered to significantly encode a variable at p ≤ 0.05. In general, a neuron’s response could be explained by multiple variables, which is likely because variables are correlated with each other, a situation referred to as multi-collinearity. We therefore adapted the “best-subset” method as a method of variable selection used in the case of multi-linear regressions (Dunn and Clark, 1987; Glantz and Slinker, 2001; Padoa-Schioppa and Assad, 2006).

#### Best-subset method

We computed for each subset of d variables the total number of neural responses explained by that subset and determined which subset explained the maximum number of responses. This was repeated for d=1, 2, 3‥ variables per subset. We determined the number of variables necessary to characterize the population when 85% of the maximum number of responses explained was reached. The best-subset method assumes that each neuron only encodes a single variable. We therefore tested for second-order encoding to determine the proportion of neurons that encoded more than one variable (Padoa-Schioppa and Assad, 2006).

#### Second order encoding

We found for each neuron the best-fit variable and its corresponding R^2^ value. To determine whether adding an additional variable to the regression led to a significantly higher R^2^ value, we computed:

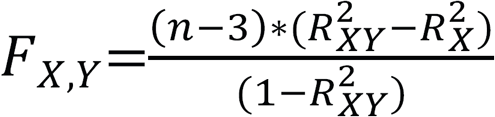

where 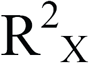 is from the original linear regression on X only, 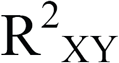 is from the bilinear regression on X and Y and n is the number of trials. F_X_,_Y_ is computed for each of the eleven possible second variables and the maximum F is found. If the corresponding p-value for the maximum F value is ≤ 0.01, we consider the neuron to significantly encode the second variable (Padoa-Schioppa and Assad, 2006). 31% of neurons significantly encoded a second task variable, which is more than expected by chance (binomial test, p < .0001). The major variables that were multiplexed at the second order were previous trial outcome (17.8%) and learning progress (26.7%), with both of these more often encoded at the second order than expected based on an equal distribution across all twelve variables (binomial test, p < .001).

